# RhoA mediates epithelial cell shape changes via mechanosensitive endocytosis

**DOI:** 10.1101/605485

**Authors:** Kate E. Cavanaugh, Michael F. Staddon, Ed Munro, Shiladitya Banerjee, Margaret L. Gardel

## Abstract

Morphogenetic movements require tight spatiotemporal control over cell-cell junction lengths. Contractile forces, acting at adherens junctions, alter cell-cell contact lengths in a cyclic fashion as a mechanical ratchet. Pulsatile RhoA activity is thought to drive ratcheting through acute periods of junction contraction followed by stabilization. Currently, we lack a mechanistic understanding of if and how RhoA activity governs junction length and subsequent cell shape within epithelia. In this study we use optogenetics to exogenously control RhoA activity in model Caco-2 epithelium. We find that at short timescales, RhoA activation drives reversible junction contraction. Sustained RhoA activity drives irreversible junction shortening but the amount of shortening saturates for a single pulse. To capture these data, we develop a vertex model modified to include strain-dependent junction length and tension remodeling. We find that, to account for experimental data, tension remodeling requires a strain-dependent threshold. Our model predicts that temporal structuring of RhoA activity allows for subsequent tension remodeling events to overcome the limited shortening within a single pulse and this is confirmed by our experimental data. We find that RhoA-mediated junction remodeling requires activities of formin and dynamin, indicating the closely inter-connected activities of contractility, E-cadherin clustering, and endocytosis. Junction length is therefore regulated by the coordinated action of RhoA-mediated contractility, membrane trafficking, and adhesion receptor remodeling. Altogether these data provide insights into the underlying molecular and biophysical mechanisms of RhoA-mediated regulation of epithelial cell shape.

## INTRODUCTION

Epithelial cell sheets dynamically restructure their shapes to sculpt higher order assemblies of tissues and organs (1–3). Constituent cells execute complex shape changes that require coordinated mechanisms to alter cell-cell junction lengths, cell surface area, and overall cell shape (4). These events are coordinated by macromolecular assemblies within the cytoskeleton that control cell force generation and junctional adhesion (1, 5). These dynamic cytoskeletal arrays underlie cell and tissue mechanics, enable them to resist deformations to maintain integrity, yet to dramatically change their shape during morphogenesis. To enable such varied and adaptive mechanical behaviors, the cytoskeleton harnesses mechanochemical feedbacks at the scale of proteins, organelles and the cell. The structures of these mechanochemical systems, and how they regulate cell physiological processes remains largely unknown.

A primary regulator of cell shape changes in epithelia is the small GTPase RhoA, which dynamically controls contractility through its downstream effectors of actin and myosin (NMII) (6, 7). Morphogenetic processes show highly dynamic RhoA activity, with pulses of active RhoA preceding the shortening of cell-cell junctions (7–9). The resulting actomyosin flows drive an oscillatory ratchet in which junctions undergo multiple rounds of contraction, stabilization, and relaxation (10–13). By spatiotemporally coordinating these pulsatile contractions within and between cells, incremental changes in cell shape collectively drive large scale tissue deformations required for apical constriction (12, 14) or convergent extension (15). While pulsatile RhoA activity is observed in diverse contexts (8, 9, 16), the significance of this temporal structure and what role it may have in junctional shortening is unknown.

Adherens junction remodeling at cell-cell contacts is essential for the maintenance of cell shapes in a variety of developmental contexts (5, 17). In Drosophila Dorsal Closure the rate of membrane removal is tension dependent to maintain a constant junctional straightness, documenting feedbacks between junctional tension and the Rab membrane trafficking pathway (18). Similarly, during Germband Extension, the Rab pathway stabilizes junction contractions during the oscillatory ratchet (19). Here, membrane tubules emanating from adherens junctions require actomyosin forces to terminate vesiculation to enforce progressive junctional shortening (19). Shear forces regulate junctional E-cadherin levels, whose clustering and subsequent internalization is also facilitated by formin and NMII activity during this cell intercalation (20, 21). Although these data suggest coupling between actomyosin contractility and membrane remodeling during junctional shortening, the underlying mechanisms remain unknown.

Here, we use optogenetic tools to modify the junctional tension in model epithelia via exogenous RhoA activation. We find that RhoA activation along the junction length leads to a rapid contraction that occurs primarily through the shortening of a few distinct micro-domains. For short activation times, this shortening is reversible such that junctions return to their original lengths after removal of the exogenous RhoA. However, as the activation time is increased, junctions shorten irreversibly, but for a single activation pulse shortening is limited. To capture these data, we introduce modifications to the existing vertex-based models for epithelial tissues that include strain-dependent remodeling of junction length and tension. In combination with experimental data, our model demonstrates that tension remodeling only occurs above a critical junctional strain. Furthermore, live cell imaging reveals that membrane remodeling and vesicular internalization occur during this RhoA-dependent junction contraction, whose stabilization at shorter lengths requires dynamin and formin activities. Finally, we show that temporally structuring the exogenous RhoA activity into distinct pulses overcomes the limited shortening. Altogether, these data provide new insights into the molecular and biophysical feedback mechanisms between RhoA activity and membrane remodeling that underlie epithelial cell shape control.

## RESULTS

### Optogenetic localization of RhoA actuates cell-cell junction shortening

To spatiotemporally control RhoA activity in model epithelia, we generated a stable Caco-2 cell line expressing the TULIP optogenetic system (22–24). TULIPs utilize the photosensitive LOVpep domain attached to a GFP-tagged transmembrane protein, Stargazin. The LOVpep’s binding partner is the prGEF complex that contains 3 components: a photorecruitable engineered tandem PDZ domain, the catalytic DH domain of the RhoA-specific guanine nucleotide exchange factor (GEF) LARG, and an mCherry fluorescent tag (Fig 1A). Blue light increases the binding affinity of the two protein complexes, recruiting the prGEF to the membrane where it locally activates RhoA (Fig. 1A) (22, 24).

**Figure 1:**
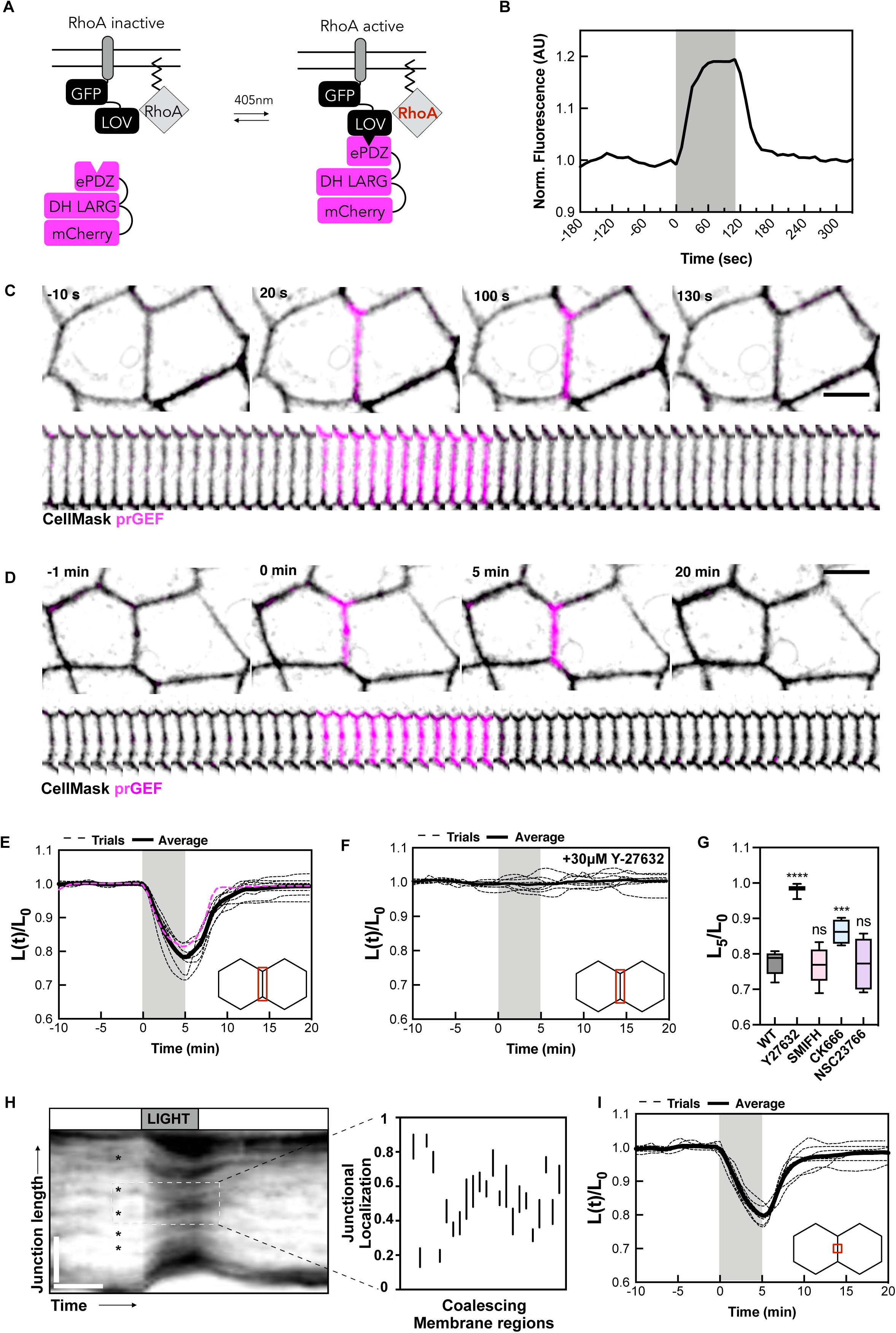
RhoA-mediated tension determines junction length at short timescales. **A.** Schematic of TULIP optogenetic system. Blue light activation causes recruitment of prGEF to the plasma membrane. **B.** Quantification of the local intensity increase of mCherry-prGEF in the junctional activation region shown in C (magenta). Activation period is indicated by the grey box. **C.** Representative images of cells undergoing junctional prGEF (magenta) recruitment. prGEF rapidly accumulates at the junction and dissociates upon light inactivation. Scale bar is 10um. **D.** Representative images of junction length changes undergoing a 5-minute activation period. Junctions undergo rapid contraction upon prGEF recruitment that is reversible upon its removal. Scale bar is 10um. **E.** Normalized junction length over time with a 5-minute activation period shows junction shortening within the activation period (grey). Individual trials are in dashed line with the average in a solid line. The magenta line represents data shown in D. **F.** Normalized junction length over time within a 5-minute activation period of cells treated with Y-27632 shows no junction length changes. **G.** Strain values (L5/L0) for WT, Y Compound, SMIFH, CK666, and nsc23766 treated cells. **H.** Representative kymograph of a junction undergoing a 5-minute activation period. Asterisks show concentrated membrane regions. White dashed box shows coalescing membrane regions. Horizontal scale bar is 6 minutes, vertical scale bar is 5um. To the right is a histogram depicting the localization of all coalescing membrane regions along all junctions undergoing a 5-minute activation. The top vertex is denoted by 1, with the bottom vertex denoted by 0. **I.** Normalized junction length over time of junctions going a 5 minute activation period with light only targeted to the center of the junctions. Junctions undergo similar contractions as full-length activation.

Cell morphologies and junction lengths in Caco-2 monolayers are incredibly stable; over 1 hour, the junction lengths change less than 3%. The remarkably static nature of the monolayer makes it particularly well suited to study junctional response to acute optogenetic stimulations, as little endogenous activities occur. Here, we target light to a desired cell-cell junction (i, Cellmask, black) (Fig. 1C) and see rapid prGEF (magenta) association (20 sec) and dissociation (30-50 sec) kinetics when the light is turned on and off, respectively (Fig. 1B&C, Video 1). Consistent with previous studies, prGEF recruitment is tightly localized to the targeted region (2, 3). The targeted junction rapidly contracts upon light activation, shortening from 13.1 um to 10.3 um over 5 min and then returning to its original length once light is removed (Fig. 1D, E pink line, Video 2). The rate and extent of junction contraction is remarkably consistent across junctions with varying initial lengths and geometries; junctions consistently contracted to 75-80% of their length at an average contraction rate of 0.047 min^-1^. To confirm that the light-actuated contraction arises from downstream effects of Rho localization, we treated the monolayer with the Rhokinase inhibitor Y-27632. Treatment with Y-27632 abolishes light-actuated junction shortening (Fig. 1F, G, Video 3).

To investigate how known actin assembly factors impact junction contraction, we treated the model epithelia with several pharmacological inhibitors. We then quantified the amount of junctional contraction, as calculated by dividing the junction length, L, at five minutes (L_5_) by its initial length at 0 minutes (L_0_). Interestingly, treatment with the pan formin inhibitor SMIFH2 and Rac inhibitor nsc23766 did not produce a significant change in initial contractile strain (Fig 1G, SFig 1). However, treatment with the Arp2/3 inhibitor CK666 significantly reduced junction contraction, suggesting that this pool of branched actin filaments is necessary for efficient junctional contraction (Fig 1G, SFig 1). These data are consistent with previous work reporting how Arp2/3 supports junctional tension at the zonula adherens (25).

Junction shortening arising from contractions along the junction length could either occur uniformly or be heterogeneously distributed. We used the variations in membrane fluorescence intensity as fiduciary markers to examine the spatial distribution of junctional movement. A line scan along the junction, observed over time, reveals how different junction portions rearrange prior to, during, and after light-mediated shortening (Fig. 1H). For a contraction that occurs uniformly along the junction length, we would expect the central portion to stay fixed and the rate of movement towards this central region to increase smoothly to the junction end. However, the kymograph data indicate that the junction does not contract uniformly. Rather, there are distinct regions that contract, indicated by converging fiduciary marks, separated by non-contractile regions, indicated by neighboring fiduciary marks that remain parallel. We identify the distinct contractile regions in junctions from the kymographs and record their size and positions along the junction length. We find that junctions typically contain 1-2 contractile regions that are several micrometers in length. In plotting a histogram of these coalescing membrane regions from the top vertex (1) to the bottom vertex (0), we found that these coalescing regions were typically found within the central portion of the junction with average normalized junctional localization to be at 0.45 (Fig. 1H). Consistent with heterogeneous junction contraction, illuminating only one third of the junction length induced similar contraction rates to that of full-length activation (Fig. 1I, Video 4).

### Extent of junction contraction saturates for a single Rho activation

Our optogenetic platform allows us to systematically vary the duration, intensity, and location of exogenous activation to determine the structure of the cellular response. To determine how junction shortening depends on the duration of prGEF junctional localization, we varied the duration of the activation pulse from 2.5 to 40 minutes. For all activation periods less than 5 minutes, junction shortening is reversible with the final length, L_f_, over initial length, L_0_, being equal to 1 (Fig 2E). However, as the activation time increases to 10 minutes, a pronounced biphasic nature of the junctional contraction dynamics is revealed. Here, the faster initial contractile phase lasts for approximately five minutes and is followed by a slower contractile phase over which junction length decreased to ~65% of its original length (Fig. 2B, Video 6). After light removal, the junction does not return to its original length. Instead, the final junction length is 20% shorter than its initial length. Similar kinetics are observed for 20-minute activation scheme (Fig. 2A, 2C, Video 5) with the slow phase showing clear saturation with a 40-minute activation time (Fig. 2D, Video 7).

**Figure 2:**
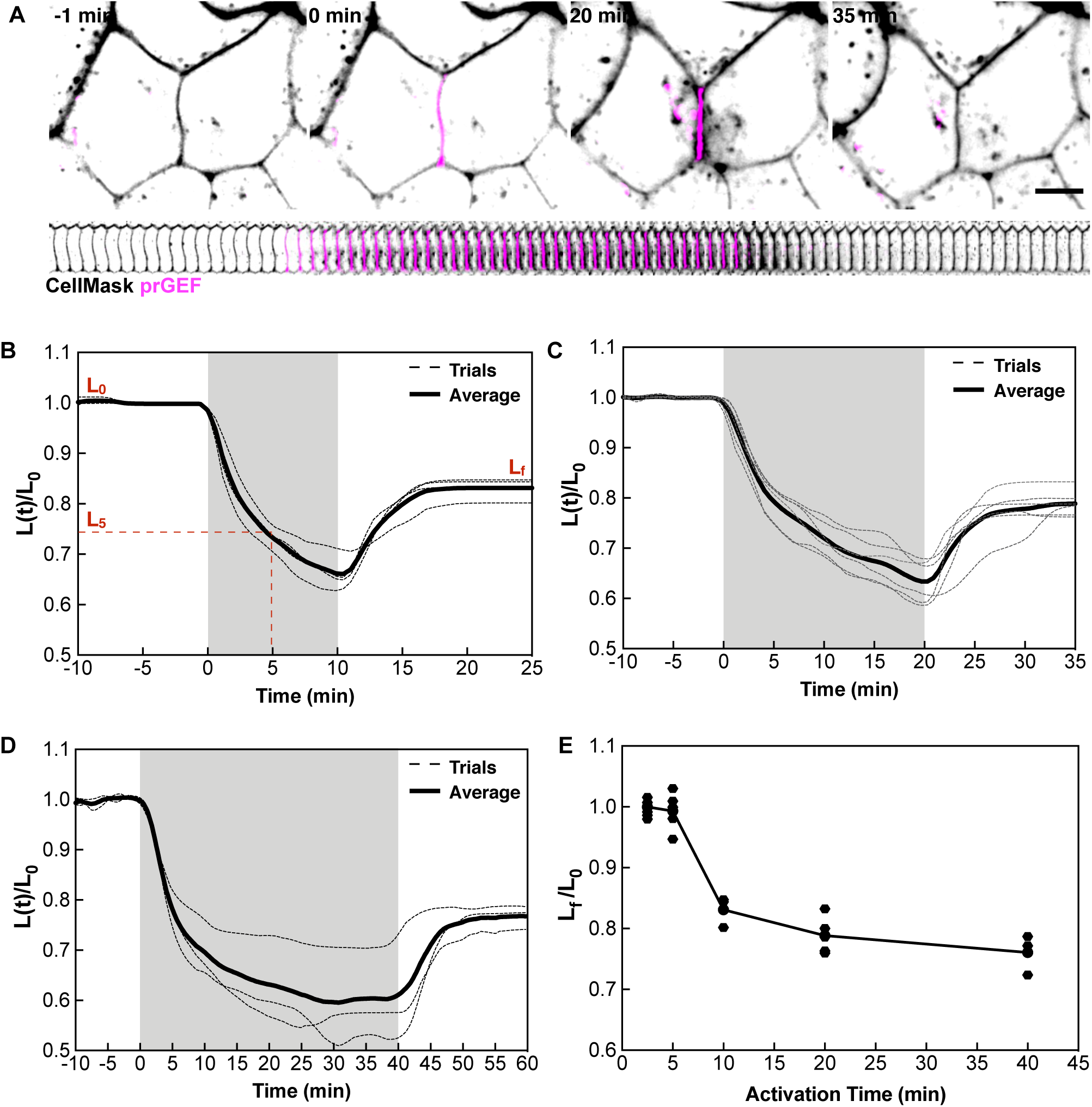
Contractility saturates at longer timescales to limit junction length changes. **A.** Representative image of a junction undergoing a 20-minute activation show similar association and dissociation kinetics as shorter activations. **B.** Normalized junction length over time for a 10-minute activation shows junction length changes to 80% of L0. **C.** Normalized junction length over time for a 20-minute activation shows junction length changes to 80% of L0. **D.** Normalized junction length over time for a 40-minute activation period shows junction length changes to 80% of L0. **D.** Plot of Lf/L0 vs Activation time shows that under 5 minutes, junction lengths are reversible, but contractility saturates at 20% reduction in junction length even at long timescales.

To explore how the activation time controls the final junction length, we plot L_f_/L_0_ as a function of activation time (Fig. 2E). This reveals that between activation times of 5-10 min, a rapid transition to permanent junction shortening occurs. Furthermore, for a single activation pulse, the shortening saturates at a 20% length reduction, even for activation times up to 40 min (Fig. 2E). Collectively these data show how the temporal structure of RhoA activation can control junctional contraction through distinct timescales, but the extent of shortening is limited.

### Strain-dependent tension remodeling captures adaptive junctional length changes

To develop a mechanistic understanding of junctional length regulation in response induced contractions, we developed a vertex-based model of epithelial monolayers. In traditional vertex-based models (26), the monolayer is modeled as a network of junctions with tension *Λ* and area compressibility *k*_*c*_. The junction lengths, *L*, are determined by the balance between junctional tension and cell elasticity (Fig. 3A). The optogenetic experiment is modeled by a step increase in junctional tension, ΔΛ, that drives length contraction and, when removed, junctional tension and lengths fully recover to their initial values (Fig. 3B, solid gray). While the vertex model captures the junction dynamics in response to short timescale activations, standard and existing variants of the vertex model(27), as well as other mechanical models of cell junctions(28–30), fail to capture the biphasic irreversible shortening observed with longer activations (Fig. S2). Thus, our experimental data are inconsistent with existing mechanical models of epithelial tissues.

**Figure 3:**
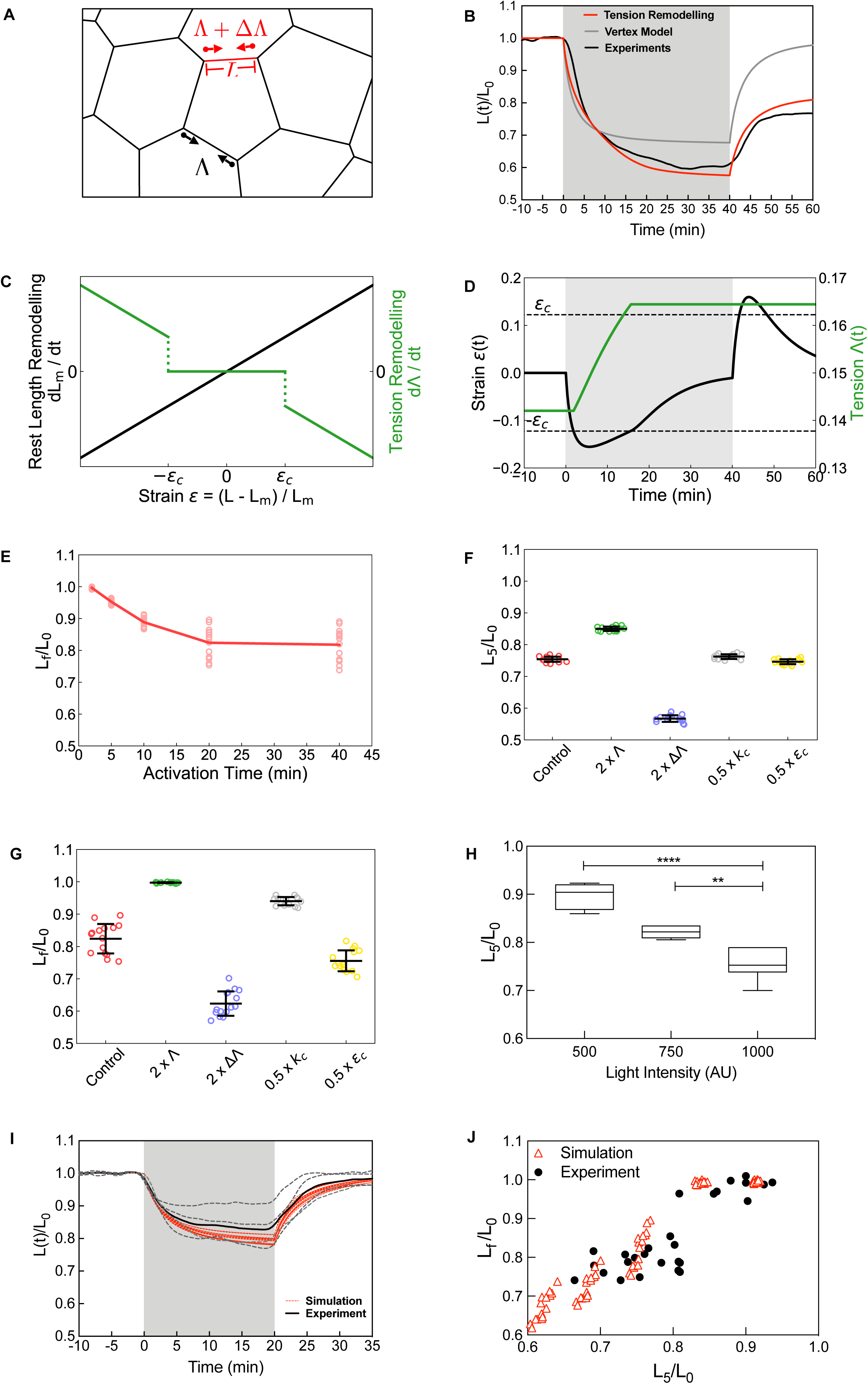
Mechanical model for mechanosensitive junction length changes. **A**, A representative image from the vertex model. Vertices are labelled by Latin symbols, and cells by Greek symbols. A line tension acts along the black edges. During an optogenetic activation, the red edge has an increased line tension along it. **B**, Normalized junction length over time with a 40-minute activation, using the vertex model with (red) and without (grey) tension remodeling compared to experiments (black). **C**, Rest length remodeling rate and tension remodeling rate as a function of edge strain. **D**, The strain and tension over time corresponding to the 40-minute activation shown in B. **E**, Final normalized junction length against activation time. **F**, Final normalized junction length after a 20-minute activation, and **G**, maximum strain over 5 minutes of activation, for different parameter perturbations. **H**, Experimental strain rate as a function of light intensity shows decreasing strain with decreasing light intensity. **I**, Normalized junction length over time using a lower strain rate, from experiments (black) and model (red). **J**, Final normalized junction length against the maximum strain over 5 minutes of activation. Data is pooled from all light intensities as all longer-timescale activations (ie. 10, 20, and 40 minute activations).

Instead, we found that both junctional length remodeling and tension remodeling are necessary to recapitulate the experimental data (Methods). In our model, each junction has a rest length that remodels at a rate to match the current junction strain, ε (Fig. 3C, black solid line). The junctional tension similarly remodels, but only when the strain is larger than a threshold value, ε_c_ (Fig 3C, green dashed line). When the exogeneous tension is applied for short times, the critical strain is not reached and, thus, junctional tension remains constant. As a result, the junction length recovers to its original value upon exogenous tension removal. However, over longer timescales and sufficiently strong ΔΛ, tension remodeling results in a permanent increase in junctional tension (Fig. 3D). Contraction stalls when the rest length remodels to the current junction length (Fig. 3D). Upon tension removal, the junction adapts to a new rest length (Fig. 3B red and black lines, 3D). By benchmarking our model parameters with experimental data (Methods) we quantitatively capture our time-dependent junction length changes over a wide range of activation periods (2.5-40 min) for a single set of parameters (Fig. S3A-E), including the changes in the final junction lengths as a function of activation time (Fig. 3E). Within this model, the initial junctional contraction is modulated by changes to base junctional tension Λ or exogeneous tension ΔΛ, while being robust to changes in other mechanical parameters. This is illustrated by plotting how L_5_/L_0_ varies as each of the parameters changes by two-fold (Fig. 3G). By contrast, the final junction length is sensitive to both junctional tension and cell compressibility *k*_*c*_, as these determine the junctional strain, as well as the critical strain required for junctional remodeling (Fig. 3F).

A key prediction of our mechanical model is that a critical strain is necessary to initiate junctional tension remodeling to enable irreversible shortening. To test this prediction, we sought to reduce the initial contraction amount by reducing the prGEF recruitment, controlled by the light intensity (Fig. 3H). When the light intensity is 1000AU, the average junction shortens on average to 20-25% of their original length after 5 minutes, with an average L_5_/L_0_ equal to 0.77 (Fig. 3H). By reducing the activation light intensity by 25% (750 AU) and 50% (500AU), the amount of initial contraction is reduced, increasing L_5_/L_0_ to 0.81 and 0.89, respectively (Fig. 3H, Video 8). When L_5_/L_0_<0.8, we find that the final length is proportional to the amount of initial contraction. However, when the initial contraction is less than 20% (L_5_/L_0_ > 0.8), we find that no junctional shortening occurs (L_f_/L_0_≈1). These experimental results have remarkable agreement with those predicted by the model when ΔΛ is varied to modify the initial contraction (Fig. 3I&J, Video 8). These findings demonstrate that a mechanosensitive junctional remodeling pathway stabilizes junction lengths upon temporal changes in Rho signaling.

### RhoA-induced junction contraction initiates membrane thickening and internalization

To gain insight into the cell biological mechanisms underlying junctional tension and length remodeling, we examined changes in the cell membrane that occur during Rho-mediated junctional shortening. We measure the membrane intensity across the junction, averaged along the junction length at several times, t=0, 5 and 20 min, during activation (Fig 4A). Initially, the line scans reveal a narrow intensity distribution of the membrane, with a full-width at half max of 0.75 μm (Fig. 4B&C) that reflect membrane localization at the cell-cell interface. During stimulated junction contraction, both the peak and width of the membrane staining increases (Fig. 4B,C), presumably reflecting the accumulation of membrane during contraction. To measure the change in membrane intensity proximal to the activated junction, we measure the average intensity in a region a distance 1.5μm away from the peak intensity and observe a dramatic increase during both the fast and slow contraction phases (Fig. 4D). Close examination of these regions reveals membrane compartments and vesicles emanating from the junction during the slow contractile phase of activation(Fig. 4E).

**Figure 4:**
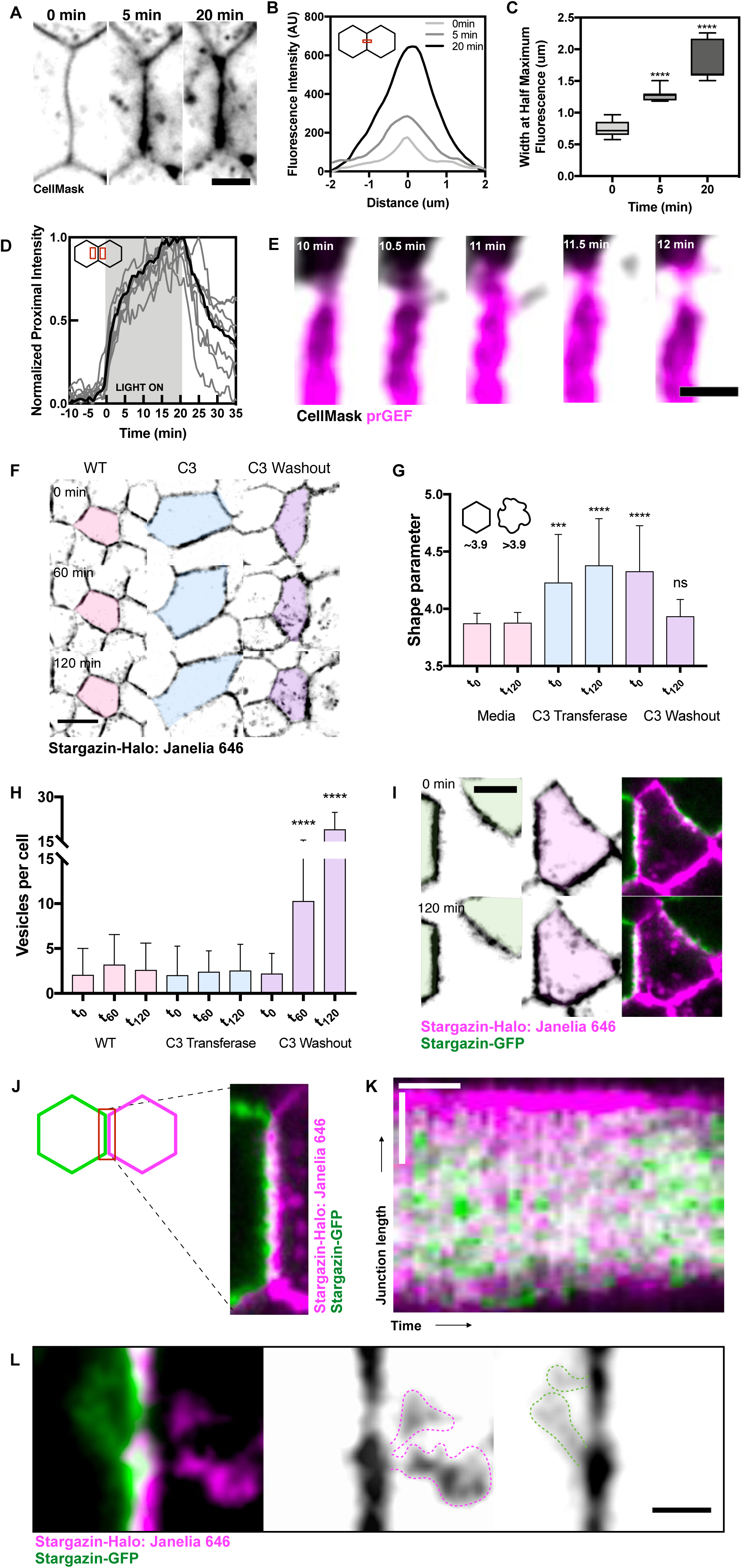
Compressive strain induces the remodeling of slackened membrane. **A.** Representative images of a junction at t0, t5, and t20. Scale bar is 5um. **B.** Representative image of fluorescence intensities of a 3um region along a junction show increasing fluorescence over time. **C.** Calculation of full width at half maximum fluorescence for all junctions shows increasing membrane thickness over time. **D.** Normalized proximal fluorescence intensities for regions alongside the activated junction shows drastic increases in fluorescence in the slow phase. **E.** Representative image of a junction in the slow phase undergoing vesiculation and compartment internalization. Scale bar is 2um. **F.** Images at 0, 60, and 120 minutes of cells in no treatment, C3 transferase treatment, or C3 transferase wash outs. Scale bar is 10um. **G.** Quantification of shape parameters from F. **H.** Quantification of vesicle internalization from F. **I.** Representative images of a mosaically labelled cell (pink) undergoing self-internalization from 0 to 120 minutes. **J.** Representative image of a mosaically labelled cell-cell junction showing heterogeneous distribution of membrane. **K.** Kymograph of junction undergoing shortening in I shows distinct membrane regions appearing during junction shortening. Horizontal scale bar is 20 minutes, vertical scale bar is 5um. **L.** Representative image showing tubular vesicles emanating off of buckled membrane regions during junction shortening. Scale bar is 2um.

To assess the role of RhoA-mediated contractility on regulating membrane internalization and cell shape changes, we monitor cell shape and membrane internalization under varying levels of RhoA activity modulated by its inhibitor, C3 transferase. In control conditions, time-lapse imaging of monolayers expressing a Stargazin-Halo membrane marker showed little cellular movement over the course of two hours (Fig 4F, Video 10 left). We quantify cell shape using a dimensionless shape parameter, defined by the ratio of the cell perimeter to the square root of the cell area (31). In control conditions, the shape parameter is 3.9, indicating a compact, hexagonal shape (Fig 4G). In this case, the average number of fluorescently labeled vesicles per cell remains low (<4) over two hours, consistent with a low amount of membrane remodeling during this time (Fig 4H).

To globally inhibit Rho activity, we incubate the monolayers in media containing C3 transferase for four hours. Rho inhibitor treatment increased the cell shape parameter to 4.5, reflecting a more elongated shape and increased cell motion (Fig. 4G, Video 10 middle). The mean number of vesicles per cells remained low (~ 2) over 2 hours. Thus, global RhoA inhibition modulates the overall cell shape, but does not modify the amount of membrane internalization that occurs over several hours.

To explore how global changes in Rho activity modulate cell shape and surface membrane reorganization, we visualize changes in cell shape and vesicle number upon washing out C3 transferase. After C3 transferase washout, the cell shape parameter decreases from 4.3 to 3.9, consistent with the increased contractility driving the transition from a more elongated to hexagonal shape (Fig 4G, Video 10 right). Strikingly, this change in cell shape is accompanied by significant increase in cytoplasmic vesicles, increasing to a mean of 10-19 vesicles per cell two hours after C3 washout (Fig 4H). Thus, we observe a significant increase in internalized membrane that occurs during Rho-mediated cell shape changes.

To further examine Rho-mediated membrane remodeling, we generated mosaic monolayers of cell lines expressing spectrally distinct membrane tags, Stargazin-Halo:Janelia-646 and Stargazin-GFP (Fig. 4J). This mosaic labeling enables the identification of the membrane source during Rho-mediated shape changes. The internalized vesicles 2 hours after C3 washout had the same fluorescence as the plasma membrane, indicating that membrane internalization all originates from the surrounding plasma membrane and does not include membrane from neighboring cells (Fig. 4I). Examining the cell-cell junction at distinctly labeled neighbors with high resolution, features along the adherens junction, including heterogeneous distribution of membrane along the junction length, can be resolved (Fig 4J). Kymographs of junctions during Rho-mediated shortening show that this heterogeneous distribution of neighboring membrane persists but varies over time (Fig 4K). This indicates that, along the adherens junction, membrane materials from one cell abruptly disappears from the junction. Examining the regions proximal to the adherens junction at these time points reveals that membrane internalization occurs via extended, tubule-like structures from concentrated membranous regions (Fig 4L). These data indicate that during Rho-mediated junction shortening, membrane at the adherens junction undergoes self-internalization within micrometer-sized domains along the junction.

### Permanent junction shortening requires dynamin-mediated endocytosis of E-cadherin clusters

It has been previously reported that adherens junction remodeling during morphogenic processes is mediated by the internalization of junctional components, including E-cadherin (20). We therefore sought to determine the relationship between RhoA-mediated membrane remodeling and junctional components. Immunofluorescence images of E-cadherin reveals it exists as distinct puncta along the lateral junction of the epithelial monolayers (Fig. 5A). Consistent with previous work (20), we find this clustering is formin-dependent as treatment with the pan formin inhibitor abolishes E-cadherin punctae (Fig. 5A).

**Figure 5:**
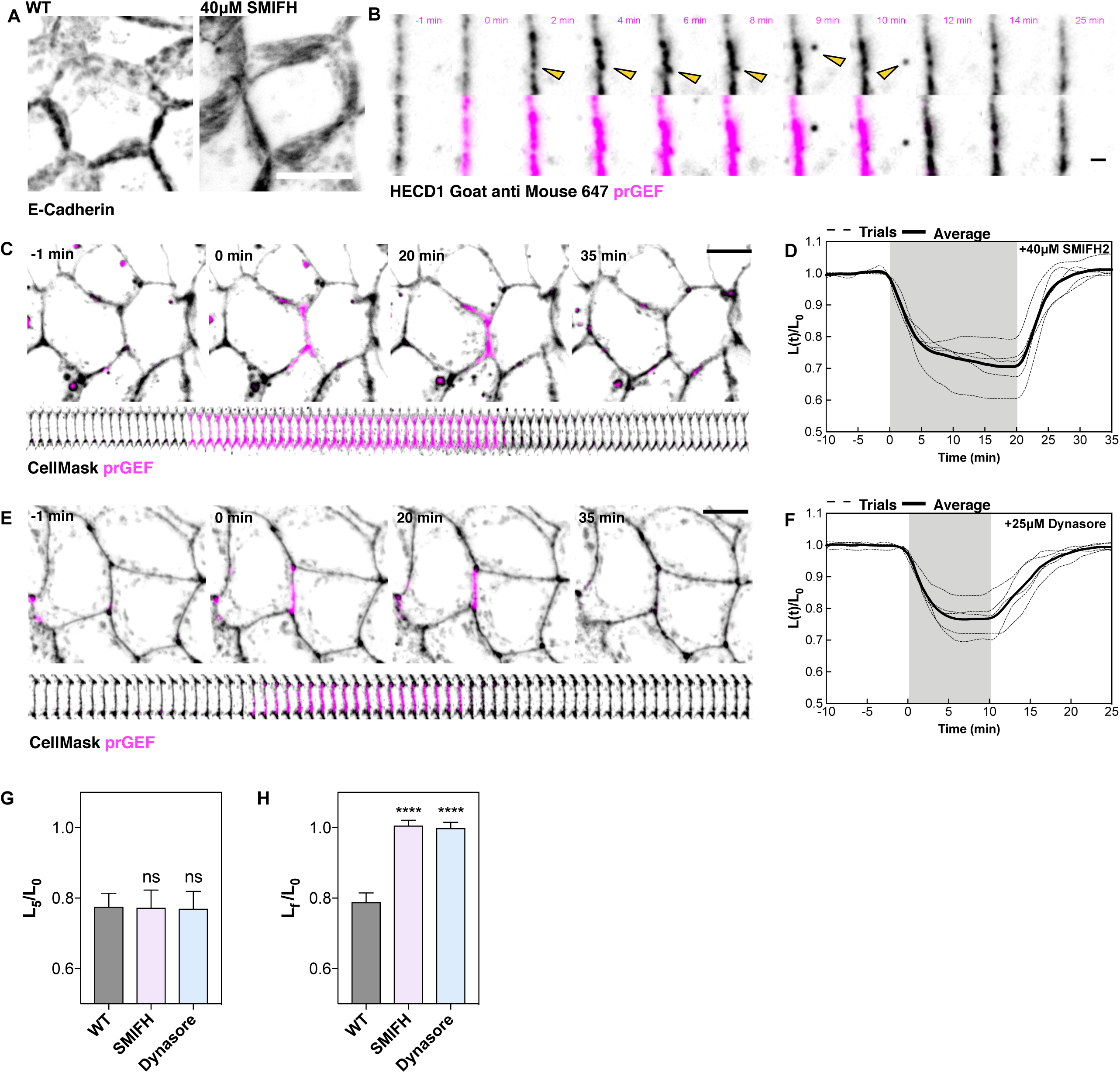
Formin-mediated E-cadherin clusters and their endocytosis are necessary for junction length changes. **A.** Representative z-stack projection of a cell stained for E-cadherin and a representative z-stack projection of a cell treated with SMIFH2 and stained for E-cadherin. Scale bar is 10um. **B.** Representative image of cells treated with HECD1 primary and GaM 647 secondary antibodies showing internalization of E-cadherin positive clusters. Scale bar is 2um. **C.** Representative images of cells treated with SMIFH2 and undergoing a 20-minute activation. Scale bar is 10um. **D.** Quantification of normalized junction lengths over time in SMIFH treated cells from C. **E.** Representative images of cells treated with Dynasore and undergoing a 10-minute activation. Scale bar is 10um. **F.** Quantification of normalized junction length over time in Dynasore treated cells from E. **G.** Quantification of strain rates for the fast phase contraction for WT, SMIFH2, and Dynasore treated cells. **H.** Quantification of the final length over the initial length for WT, SMIFH2, and Dynasore treated cells.

To determine how E-cadherin puncta are impacted by exogenous RhoA activation, we visualized E-cadherin in live cells by incubation with fluorescent conjugates of HECD1 antibody to preserve trans-cadherin interactions while labeling endogenous E-cadherins (32). This labelling revealed similar punctate pattern of E-cadherin along the junction and this treatment did not induce any junction length changes or endocytic events prior to light activation (Fig. S1C). Upon light stimulation, similar junctional shortening (Fig. S1, Video 11) and motion of E-cadherin puncta are observed (Fig. 5B). In particular, E-cadherin puncta coalesce together, intensify, and are then are internalized over the course of a 10-minute exogenous Rho stimulation (Fig. 5B, Video 12). Thus, internalization of E-cadherin clusters occurs during junctional shortening via exogenous RhoA.

Regulation of endocytosis can occur either within the clustering or vesiculation phases (20). We therefore sought to elucidate the molecular regulation of observed membrane remodeling and internalization. To test the role of E-cadherin clustering, we treated the monolayers with SMIFH2 and stimulated the junction will full intensity light for 20 minutes (Fig. 5C, D). As described earlier, SMIFH2 treatment had no impact on the initial contraction rate and the extent of junctional shortening after 5 minutes is the same as control conditions (Fig 5C&D, G). However, after light is removed after 20 minutes, the junction recoils back to its original length and no permanent length reduction is observed (Fig. 5D, H, Video 13). To explore the mechanism of vesiculation, we treated the monolayers with Dynasore, a pharmacological inhibitor of dynamin. Phenocopying SMIFH treatment, the monolayer treatment with Dynasore did not alter the initial fast contractile phase (Fig 5E&F) and no permanent reduction in junction length is observed after blue light removal (Fig 5F, H, Video 14). Thus, the permanent junction shortening that occurs after a pulse of heightened RhoA requires formin-mediated E-cadherin clustering and dynamin-mediated endocytosis.

### Pulsatile RhoA Enables Junctional Ratcheting

Our experimental and modeling data show that junction shortening from a single step increase in RhoA activity saturates at ~20%, even for longer activation times (Fig. 3E). Within the model, the underlying mechanism for junction length regulation originates from thresholded tension remodeling (Fig. 3C). With a step increase in exogeneous tension ΔΛ (Fig. 3A), the junction tension transiently remodels until the junctional strain falls below the threshold ε_c_ (Fig. 3D). This increased tension results in a reduced junction length after the removal of the exogenous tension. Subsequently, the junctional strain increases transiently but does not increase sufficiently high to drive tension remodeling and reduces under the non thresholded-length remodeling. Thus, for a single activation, junctional shortening is limited by the amount of tension remodeling that occurs upon the initial onset of increased contraction.

However, this framework suggests that a subsequent exogenous pulse could overcome the saturation in junction shortening, provided ΔΛ is sufficiently large to initiate a second phase of tension remodeling. To explore this, we performed simulations with two pulses of ΔΛ separated by varying amounts of “rest” time when the exogenous tension ΔΛ is removed (Fig. 6A). During this rest period, the tension removal results in junction length extension (Fig. 6A) that is relieved by length remodeling that relaxes the strain (Fig. 6B). Consequently, the second exogeneous tension pulse results in a sufficient contractile strain to drive a second period of tension remodeling (Fig. 6B). The extent of tension remodeling that occurs in the second pulse depends on the strain at the end of the relaxation time, ε_r_ (Fig. 6B). Consequently, the amount of additional length contraction that occurs between the second and first pulse (Fig. 6A) is proportional to ε_r_ (Fig. 6C). We then explored the state space of the amount of final junction shortening that is observed for varying total duration of contraction time of two identical pulses that are separated by varying rest times (Fig. 6D). Consistent with our previous results, when the rest time is short, junction contraction saturates even for longer activation times. However, as the rest time period increases, temporally structuring the contraction into two distinct pulses enables further junction shortening for the same amount contraction duration. Thus, our model for mechanosensitive kinetics of junction remodeling leads to the prediction that frequency modulation of Rho GTPase activity will have significant impact on junction shortening than amplitude modulation.

**Figure 6:**
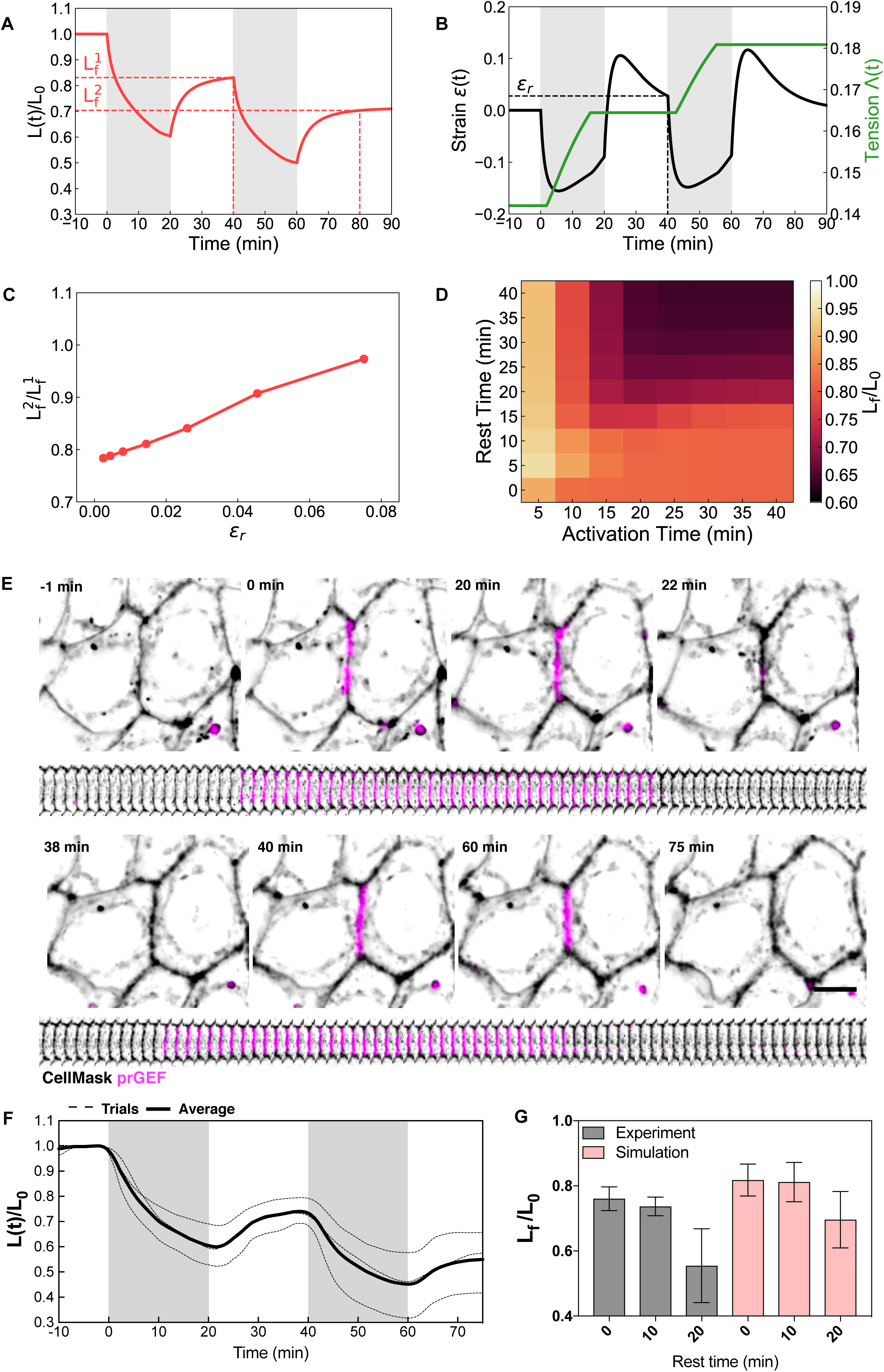
Pulsatile RhoA Enables Junctional Ratcheting. **A**, Normalized junction length, and **B**, junction strain and tension, during two 20-minute activations with a 20-minute rest. **C**, Ratio of junction length after second activation over length after first activation, against junction strain after the first activation. Each point is the average value from different rest times. **D**, Final junction length after two activations as a function of rest time and activation time. **E**, Representative image of a junction undergoing two 20 minute activations with a 20 minute relaxation period in between. **F.** Normalized junction length of junctions undergoing two 20 minute activations with a 20 minute rest in between activations. **G.** Final normalized junction length against rest time in experiments (grey) and simulations (pink). Error bars indicate standard deviation.

To test the predictions of our model, we modify the temporal pattern of exogenous RhoA activation and compare the total length change induced by a single 40-minute light activation (Fig. 2D) to that of two 20-minute pulses separated by either 10 minutes or 20 minutes rest. As described previously, a single 40-minute pulse shortens the junction length by 20%. The dual pulses separated by 10 minutes shows a similar extent of contraction (Fig. 6G). However, when the relaxation time is increased to 20 minutes, the junction shortens to 40-50% of the original length (Fig 6F, G, Video 15), in agreement with our model predictions.

## DISCUSSION

In this study, we have utilized optogenetics to investigate Rho-dependent changes in cell shape in model epithelia. Our optogenetic system has allowed us to uncover a homeostatic mechanism of epithelial tissue in response to RhoA-mediated strain. Here, small junction length changes are reversible such that tissue shape homeostasis is maintained. In this way, fluctuations of RhoA are buffered, essentially preserving epithelial architecture to prevent large scale cellular or tissue shape changes under short time-scale fluctuations in RhoA. Sufficient RhoA activity for sufficiently long times triggers junctional tension remodeling that drives irreversible junction shortening. Thus, our data suggest that cell shape changes within epithelial tissues depend more on the temporal structure of RhoA activity rather than its amplitude.

These data have serious implications for the validity of commonly used cell-based mechanical models of epithelial tissues (27–30), which treat junction lengths solely as a function of junction tension and cell elasticity. While standard and existing variants of vertex-based models capture reversibility of junction length changes upon short-time scale contraction, they are inadequate to describe irreversible biphasic junction shortening or ratcheting behavior for repeated longer time-scale contractions. As these models failed to recapitulate our experimental data, we developed a new vertex-based model where permanent junction length changes are triggered by strain-sensitive tension remodeling in cell junctions. Our model for junction tension remodeling suggests that the junctions have an inherent strain threshold that buffers small changes in contractility. Longer periods of strain changes separated by sufficient periods of strain relaxation can then direct major morphogenetic movements in epithelial tissues.

We find that membrane remodeling is critical to Rho-mediated junctional shortening. This is in line with recent work showing the role of the Rab trafficking pathway in controlling epithelial cell shape changes (18, 19, 33). Our data show that junctional RhoA activation drives heterogeneous contraction of distinct junctional regions. This compression thickens the membrane to isolate distinct microdomain buckles that initiate remodeling of the slackened regions through a mechanism that is formin and dynamin dependent. These mechanisms regulating junction length likely represent a conserved morphogenetic mechanism of general junction removal. Internalization of slackened membrane has been seen in other developmental contexts such as Drosophila Dorsal closure, in which junction length is maintained due to a straightness-dependent junctional membrane removal mechanism (18). Altogether, these data propose a general model whereby compressive strain induces membrane buckles that either initiate or control the rate of endocytosis for cell shape stabilization. It will be of interest to explore any feedback mechanisms between RhoA and the general endocytic machinery. Already it has been shown that Dia signaling, downstream of RhoGEF2, can induce ectopic clathrin recruitment at cell junctions during major morphogenetic changes (20).

We find that two distinct pools of actin are necessary for Rho-mediated junction contraction and shortening. Our data suggests that Arp2/3 plays a role in the initial contraction but is dispensable for permanent junction shortening. This finding stands in contrast with previous work documenting junction length changes being dependent on Arp2/3-mediated vesicular scission (20). Instead, we find that junctional remodeling requires both formin and dynamin. Formin dependent E-cadherin clustering is required for the initiation of internalization (20). Dynamin is essential for several endocytic pathways (34) and has been also shown to organize contractile actomyosin (35). We hypothesize that Rho-mediated contractility induces the coalescence of E-cadherin clusters at points of slackened regions. Formin activation therefore provides contractile actomyosin patches that concentrates E-cadherin clusters to facilitate their endocytosis. Without formin or dynamin, tubulation may occur, but the junction becomes reversibly deformable to prevent material internalization and subsequent junctional remodeling.

That E-cadherins are internalized during polarized compressive strain is in line with recent work reporting how differences in the subcellular origin of contractile forces affects junctional E-cadherin endocytosis (21). For instance, polarized junctional NMII generates shear stress that destabilizes and dissociates E-cadherin trans-interactions for their subsequent internalization via heterogeneous load distribution (21). This is in line with our work documenting punctate patterns of local E-cadherins that we believe generate the distinct contractile units along the junction that drives junction compressibility. We propose that these contractile units generate regions of slackened membrane which act as endocytic hubs, and these regions are tightly localized to the center of the junction. Our data further proposes that the center of the junction is the main proponent of shear stress that is capable of remodeling adhesive complexes across cell membranes.

Our optogenetic system, in combination with a new junction-scale model for epithelium, has allowed us to dissect how local, pulsatile RhoA can be used to control epithelial junctional length and tension. In vivo, junction contraction is cyclical as an oscillatory ratchet driven by RhoA pulses (8, 9). Post contraction, there are brief periods of junction expansion before the next round of contractions (13). Recent work has shown that apical constriction relies on the cycling of Rho and subsequent pulsatile contractions, inhibition of which completely halts tissue scale morphogenesis (16). While the biological function of this ratcheting behavior has remained elusive, our data highlight a function for temporally coordinated Rho pulses. Indeed, at longer timescales, we find that contractility saturates for a period of sustained RhoA activation. Instead, we find that simulating ratcheting produces multiple critical strain events and successfully produces junction length changes past this 80% value. We therefore conclude that pulsatile contractions are likely not a by-product of actomyosin network dynamics, but rather serve an essential function in coordinating cell and tissue-scale morphogenetic movements.

## Materials and methods

### Plasmids and Generation of Cell lines

Lentiviral vectors described here were generated with the aid of SnapGene Software (GSL Biotech LLC). The optogenetic membrane tether consisting of Stargazin-GFP-LOVpep and prGEF constructs were constructed by PCR amplification of the region of interest and inserting it into a lentiviral vector, pWPT-GFP (12255; Addgene) using restriction sites BamHI and SalI. This created the lentiviral constructs WPT-Stargazin-GFP-LOVpep and WPT-mCherry-prGEF. pWPT-Stargazin-Halo was constructed by inserted a PCR amplified a Halo tag region into the the pWPT-Stargazin-GFP-LOVpep backbone isolated by PCR amplification. Stargazin-GFP was constructed by PCR amplification and insertion into pWPT-GFP using restriction sites BamHI and SalI.

Lentivirus was produced using 293T cells (a gift from G. Green, University of Chicago, Chicago, IL) using Fugene 6 transfection reagent (Roche) to transfect the lentiviral vectors, a pHR1-8.2-delta-R packaging plasmid, and VSV-G pseudo typing plasmid (gifts from M. Rosner, University of Chicago, Chicago, IL). Viral supernatant was collected, filtered, and incubated with target Caco-2 cells for 24hr in the presence of 8ug/ml polybrene (EMD Millipore). After viral transfection, cells were sorted by fluorescence (UChicago Flow Cytometry Core) and screened for optimal expression of constructs by fluorescence microscopy.

### Collagen gell preparation

Collagen gels were prepared by mixing Rat tail collagen 1 (Corning) with collagen polymerizing reagents to a final concentration of 2mg/ml. Collagen polymerizing agents were prepared with media mixed with 1M Hepes and 7.5% NaHCO3 for a final ratio of 1:50 and 1:23.5, respectively. Four-well chambers (Ibidi) were coated with 30ul of 2mg/ml collagen solution and polymerized in the incubator for 10min before plating cells.

### Microscopy

Cells were imaged on an inverted Nikon Ti-E (Nikon, Melville, NY) with a Yokogawa CSU-X confocal scanning head (Yokogawa Electric, Tokyo, Japan) and laser merge model with 491, 561, and 642nm laser lines (Spectral Applied Research, Ontario, Canada). Images were collected on a Zyla 4.2 sCMOS Camera (Andor, Belfast, UK). Optogenetic recruitment utilized a Mosaic digital micromirror device (Andor) coupled to a 405nm laser. A 60x 1.49 NA ApoTIRF oil immersion objective (Nikon) or a 60x 1.2 Plan Apo water (Nikon) objective was used to collect images. MetaMorph Automation and Image Analysis Software (Molecular Devices, Sunnvyale, CA) controlled all hardware.

### Live-cell imaging

To ensure a mature and polarized epithelial monolayer, Caco-2 cells were plated at a confluent density on a collagen gel within an Ibidi chamber and cultured for two days prior to any experiments. Ibidi chambers were then placed in culture media supplemented with 10mM Hepes and maintained at 37°C or maintained with a stage incubator for temperature, humidity, and CO2 control (Chamlide TC and FC-5N; Quorum Technologies). For the stage incubator, the stage adapter, stage cover, and objective were maintained at 37C, whereas humidified 5% CO2 was maintained at 50°C at its source to prevent condensation within its tubing.

For optogenetic experiments, cell-cell junctions were labeled with CellMask Deep Red plasma membrane stain (Molecular Probes, Life Technologies). Cells were imaged in the 561 and 647 channel every 35 seconds. The first 10min was used document average junction length fluctuations and the last 15 minutes to document junction recovery. For analysis of the timescales of RhoA activation, the activation period was 2.5, 5, 10, 20, or 40 minutes. During the stated activation period, a local region was drawn around the cell-cell junction in MetaMorph and illuminated by the 405nm laser for 1000ms immediately before the acquisition of each image. Activated regions were adjusted in real time to isolate the prGEF at contracting junctions. Unless otherwise stated, laser power was set at 1000AU. For determining junction strain rates as a function of laser intensity analysis, laser power was reduced to 750 or 500AU.

Mosaic labeling of cells was performed by mixing the two cell lines, Stargazin-Halo and Stargazin-GFP, at least a day before imaging and grown to ensure a confluent and polarized monolayer. For visualization, Stargazin-Halo cells were conjugated with the Halo ligand, JaneliaFluor 646 (a kind gift from Luke Lavis, HHMI Janelia Research Campus, Ashburn, VA).

### Drug treatments

For optogenetic experiments, cells were treated with stated drug for at least one hour before any optogenetic activations and imaging. Treatments here were 30uM Y-27632 (Sigma), 25uM Dynasore (Tocris), 40uM SMIFH2 (a gift from D. Kovar, University of Chicago, Chicago IL), 100uM CK666 (Calbiochem), or 1:1500 dilution of HECD1 antibody (Abcam) and Alexa Fluor Goat anti Mouse 647 (Invitrogen). For C3 transferase washout experiments, cells were incubated with 1ug/ul C3 transferase (Cytoskeleton, Inc.) diluted in serum-free media for 4 hours, washed with PBS, and then replaced with fresh media.

### Image analysis

For junction length analysis, junction lengths were measured manually in each frame using the free hand line tool in FIJI software. Strain and strain rates were calculated from this data using custom R scripts. Junction width analysis was done by taking a junctional region of 3um by 3um and measuring fluorescence intensities with the FIJI intensity analysis tool. Full width by half maximum was calculated by hand in Excel. Proximal fluorescence intensity analysis was done by taking a region proximal to the junction outside of the junctional regions and also measured using the FIJI intensity analysis tool. Localization of microdomains was calculated by measuring the distance of concentrated membrane regions from each vertex using the FIJI line tool. Vesicle number was calculated using FIJI by thresholding and segmenting the image to create a mask of vesicles within the cell perimeter, which was overlaid onto the channel for segmentation and measurement analysis in FIJI. Cell perimeter and area was calculated manually by tracing cell junctions with the line tool in FIJI. Shape parameter was then calculated in Excel.

### Immunofluorescence Staining

Cells were seeded onto collagen gels in Lab Tek II Chamber Slides (Thermo Fisher Scientific) and allowed to form a polarized, mature monolayer before fixing. Cells were fixed in a solution of 4% paraformaldehyde with 0.1% Triton X-100 in PBS solution (Corning). Cells were permeabilized in 0.5% Triton X-100 for 10 min and blocked with 2.5% BSA and 0.1% Triton X-100 in PBS for 1 hr. Cells were incubated with mouse HECD1 antibody (Abcam) at 1:300 in blocking solution overnight at 4°C and washed three times in 0.1% Triton X-100 for 20 min each, and then placed in secondary antibody Alexa Fluor Goat anti Mouse 647 (Invitrogen) in blocking solution for 1 hr. After another three 20-min washes with 0.1% Triton X-100, slide chambers were removed. Samples were prepared with 20 *µ*l ProLong Gold (Thermo Fisher Scientific) per well and sealed with glass coverslips. Slides were allowed to dry, sealed with nail polish, and stored at 4°C.

### Quantification and statistical analysis

Statistical significance was determined under specific experimental conditions and was established during a two-tailed Student t-test in Prism (Graphpad).

### Cell-based model for epithelium with variable tension and junctional remodeling

#### Vertex model for epithelial mechanics

Each cell is modelled by a two-dimensional polygon, with edges representing cell-cell junctions, and vertices three-way junctions. The mechanical energy for the tissue is given by(26):

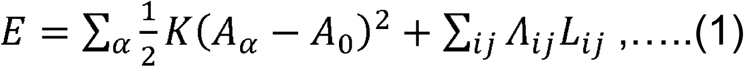

where the first term represents the area elasticity of each cell, *A*_*α*_ is the area of cell *α, A*_0_ is the preferred area, and *K* is the areal elastic modulus. The second term is the interfacial energy resulting from cortical tension and cell-cell adhesion, where *L*_*ij*_ is the length of edge *ij* connecting vertices *i* and *j*, and *Λ*_*ij*_ is the tension on that edge.

The net mechanical force acting on vertex *i* with position ***x***_*i*_ is given by ***F***_*i*_ = –∂*E / ∂****x***_*i*_. Assuming overdamped dynamics, the equation of motion for the vertex *i* is:

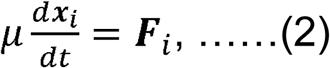

where *μ* is an effective friction coefficient. Prior to RhoA induced contractions, all cell-cell junctions are assumed to carry uniform tension, *Λ*_*ij*_ = Λ. Under applied mechanical strain both tension and junctional lengths can remodel as described below.

#### Modeling RhoA induced contractions

To model the mechanical effect of optogenetic activation of RhoA on cell junctions, we increase the tension on the activated edge *ij*, by an amount *ΔΛ = Γ*_*opt*_*L*_*ij*_, where *Γ*_*opt*_ is applied contractile force per unit length, and *L*_*ij*_ is the length of the edge. The increase in tension is assumed to be proportional to the edge length, so as to produce a strain that is independent of the initial edge length (SFig X1D), as observed in our experiments. Having a constant strain, *ΔΛ = Λ*_*opt*_, applies a higher strain on shorter edges.

#### Mechanosensitive junctional remodeling

Each cell-cell junction has a rest length, 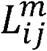, which is the length at which the junctional elastic strain is zero and the tension is constant. Prior to RhoA induced contraction of edge 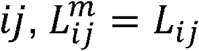. As the junctional edges contract or expand due to applied forces, the rest length remodels over time to adjust to the current length of the edge (28):

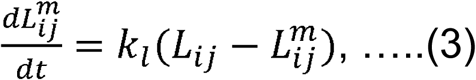

where *k*_*l*_ is the rate of rest length remodeling. When the e ge is stretched, or compressed, above a critical strain, *ϵ*_*c*_, the edge tension begins to remodel:

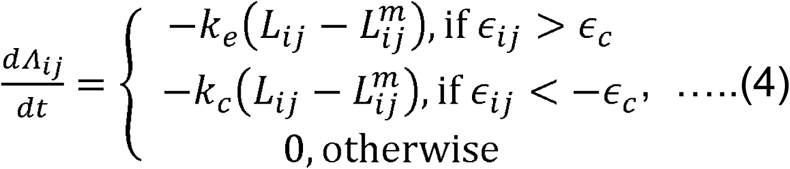

where *k*_*e*_ the rate of tension remodeling during extension, *k*_*c*_ the rate of tension remodeling during compression, and 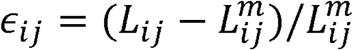 is the strain on the edge. Since cell membrane can buckle easily under compression, and stiffen under extension, we allow the remodeling rates to be different during extension and compression.

#### Short timescale contraction

If the contraction is induced for a short period such that *ϵ* > – *ϵ*_*c*_, then the edge tension does not remodel and remains constant at Λ(Eq. 4). The edge length contracts by an amount proportional to *Γ*_*opt*_, and the rest length remodels at a rate *k*_*l*_ to approach the current length (Eq. 3). As the applied contraction is turned off, *Γ*_*opt*_ = 0, the edge length fully recovers to its pre-stimulus value (Fig. 3) determined by the balance between tension (*Λ*) and elasticity.

#### Long timescale contraction

During long timescale contraction of an edge, the edge gains a permanent increase in tension due to tension remodeling (*ϵ*< -*ϵ*_*c*_) As a result, the balance between edge tension and elasticity is altered, leading to a shorter final length in the steady state after the contraction is turned off (Fig. 3).

#### Ratcheting

Since we observe that the strain on an edge is independent of the initial edge length, applying a second activation to an already-shortened junction should lead to further contraction, even in the case of a long, saturated contraction. During the first contraction, at long times the strain rate slows down and the rest length remodels and the strain falls below the critical strain, stopping tension remodeling and eventually leading to a saturation of the contraction. After activation the edge recoils to a new length. A second activation leads to a fast contraction, which strains the edge above the critical strain, and so the tension continues to increase, leading to a further decrease in junction length. Thus, applying repeated contractions can repeatedly shorten the junction.

### Different model limits

A. *Vertex model with constant tension and constant rest length* 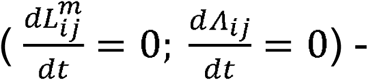 Using the traditional vertex model, we find that, regardless on contraction time, junctions always return to their initial length after contraction. Other common variants on the vertex model also fail to capture the change in length of the edges, such as including an energy term proportional to square of the perimeter) (SFig. 2A). The stable state of the junction is such that forces balance. Since forces don’t change during a contraction, the stable state remains unchanged and the junction returns to its initial length. This model is equivalent to setting tension remodeling to zero.
B. *Vertex model with constant rest length and tension remodeling 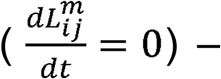* With no rest length remodelling, an activation would shorten the edge and increase the edge tension. After the activation, the new steady state length is shorter than the rest length, leading to a further increase in tension and an even shorter rest length. This positive feedback loop would lead to the collapse of any junction after an activation (SFig. 2B).
C. *No critical strain for tension remodeling* (*ϵ*_c_=0) - With *ϵ*_c_, after activations the edges slowly recover to their original length. With no critical strain, the tension increases during and activation, but immediately after activation it begins to decrease. As a result, the junction extends, further decreasing the tension until it returns to its original length. In particular, the recoil continues for a long time when compared to experiments (SFig. 2C). With a critical strain, the tension stops decreasing and a new steady state is reached.
D. *Constant applied tension* – Increasing the tension by a constant amount during an activation, Λ_*opt*_ instead of Γ_*opt*_*L*, shrinks edges at a constant speed, independent of initial length, and so shorter edges experience a much higher strain rate than longer edges (SFig. 2D). In contrast, using a tension proportional to the edge length leads to a strain rate independent of edge length, as in experiments.

### Model Parameters

Model parameters are fit to the experimental data by minimizing the error between simulated and experimental data, for 2, 5, 10, 20, and 40-minute activations. We minimize the mean square error between individual simulated lengths and mean experimental lengths during an activation during and after the activation, averaged over the different activation times (SFig. X):

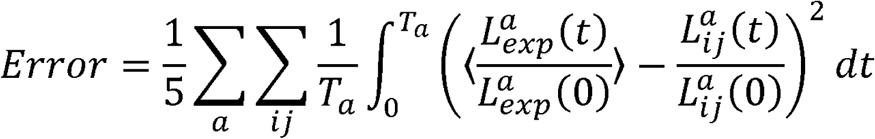

where *a*=2,5,10,20,40indicates the activation time, *ij*, indicates the simulated edge being activated, *T*_*a*_ is the total time recorded in experiments for activation *a*, and 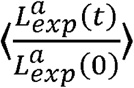 is the normalized length in experiments averaged over all activations. We then use the Nelder-Mead algorithm implemented in scipy to minimize the error, giving the parameters in the table below.

We nondimensionalize our parameters by rescaling length by 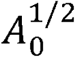 and force by 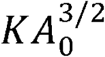, so that the nondimensional energy becomes

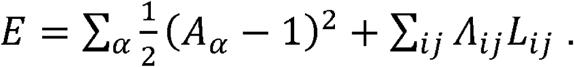

| Parameter | Symbol | Value |
| --- | --- | --- |
| Default edge tension | $\Lambda$ | 0.142 |
| Rest length remodeling rate | $k_l$ | 0.159 / min |
| Contraction tension remodeling rate | $k_c$ | 0.0203 / min |
| Extension tension remodeling rate | $k_e$ | 0.000 / min |
| Critical strain | $\epsilon_c$ | 0.122 |
| Optogenetic contractility | $\Gamma_{opt}$ | 0.0846 |
| Viscosity | $\mu$ | 1.132 mins |

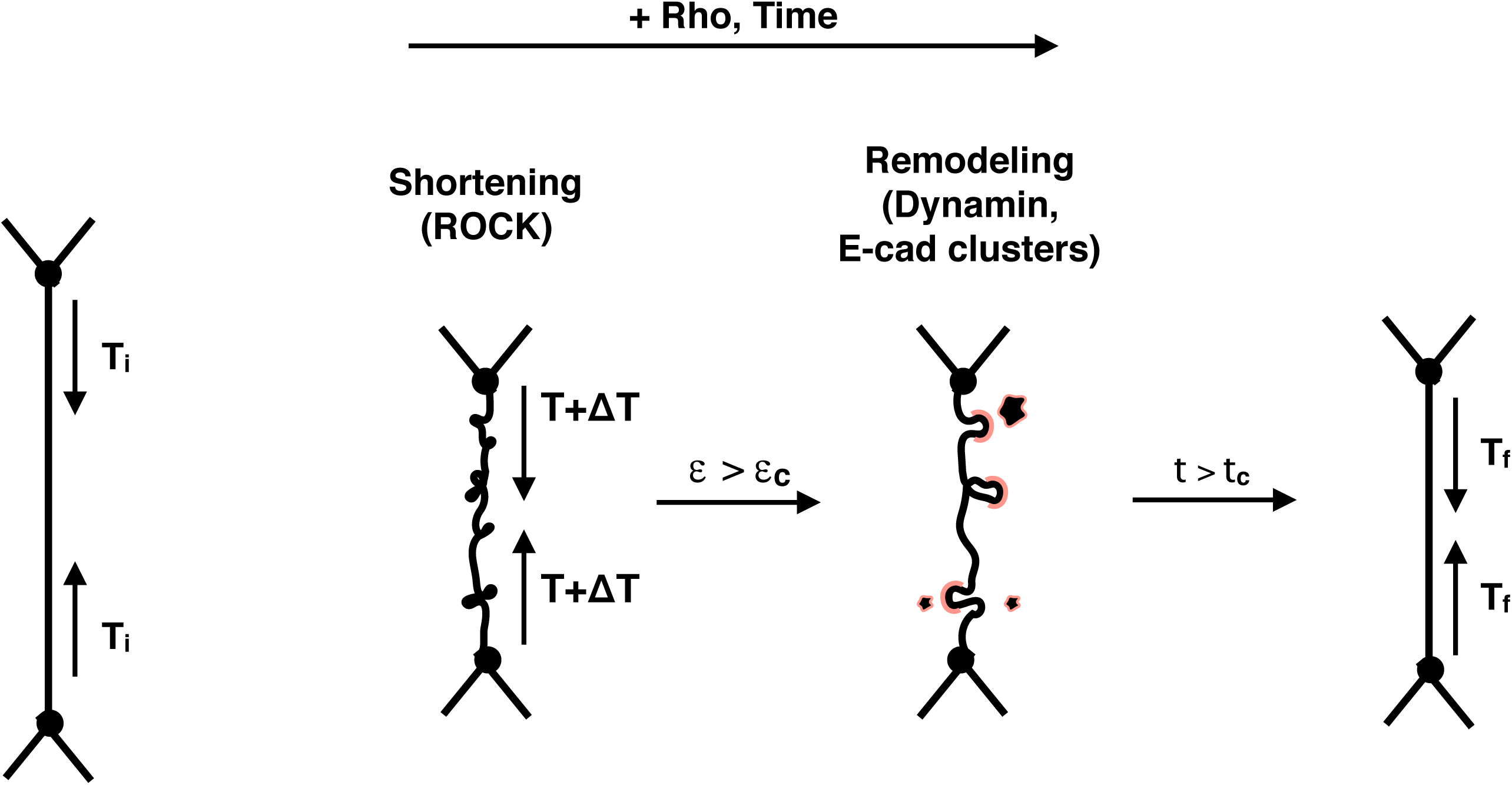

## Acknowledgements

KEC acknowledges an HHMI Gilliam Fellowship, National Academies of Sciences Ford Foundation Fellowship, and NIH training grant GM007183. MLG acknowledges funding from NIH RO1 GM104032. MFS is supported by an EPSRC funded PhD studentship. SB acknowledges funding from the Royal Society URF/R1/180187. EM acknowledges funding from NIH RO1 HD099931.

## VIDEO CAPTIONS

Video 1: Light-mediated recruitment of prGEF is rapid and reversible at cell-cell junctions (from Figure 1B, C). Zoomed in image of two Caco-2 cells during local activation of RhoA at their connecting cell-cell junction. Blue light activation period is 100 seconds. Scale bar is 10um.

Video 2: Light-mediated recruitment of prGEF causes a rapid and reversible junction contraction (from Figure 1D). Zoomed in image of two Caco-2 cells during a 5 minute RhoA activation at their cell-cell junction. The junction undergoes a contraction that is reversible upon RhoA removal. Scale bar is 10um.

Video 3: Y-27632 treatment abolishes prGEF-mediated junctional contraction (from Figure 1F). Zoomed in image of two Caco-2 cells during a 5 minute RhoA activation at their cell-cell junction post Y-27632 treatment. Junctions fail to undergo contractions. Scale bar is 10um.

Video 4: Light-mediated recruitment of prGEF to the center of the junction causes a rapid and reversible junction contraction (from Figure 1H). Zoomed in image of two Caco-2 cells during a 5 minute RhoA activation at the center of the cell-cell junction. The junction undergoes a contraction similar to full length junctional activation. Scale bar is 10um.

Video 5: Light-mediated recruitment of prGEF causes junction length changes after 20 minutes of activation (from Figure 2A). Zoomed in image of two Caco-2 cells during a 20 minute RhoA activation at their cell-cell junction. The junction undergoes a 20% length change upon light inactivation. Scale bar is 10um.

Video 6: Light-mediated recruitment of prGEF causes junction length changes after 10 minutes of activation (from Figure 2B). Zoomed in image of two Caco-2 cells during a 10 minute RhoA activation at their cell-cell junction. The junction undergoes a 20% reduction in junction length upon light inactivation. Scale bar is 10um.

Video 7: Light-mediated recruitment of prGEF causes junction length changes after 40 minutes of activation (From Figure 2D). Zoomed in image of two Caco-2 cells during a 40 minute RhoA activation at their cell-cell junction. The junction undergoes a 20% reduction in junction length upon light inactivation. Scale bar is 10um.

Video 8: Activation of a single junction under 3 different light intensities (From Figure 3H). Zoomed in image of two Caco-2 cells during 3 pulses of light at 1000, 750, and 500 AU. The junctional strain decreases with decreasing light intensity. Scale bar is 10um.

Video 9: Light-mediated recruitment of prGEF at 500au causes reversible junction shortening (from Figure 3G). Zoomed in image of two Caco-2 cells during a 20 minute RhoA activation at half the light intensity. The junction undergoes a 10% reduction in junction length before returning to its original length upon light inactivation. Scale bar is 10um.

Video 10: Heightened RhoA levels induce cellular-level shape changes (from Figure 4F). Zoomed in image of Caco-2 cells expressing a membrane label, Stargazin-Halo conjugated with Janelia Fluor 646. Cells are treated with media (left), C3 transferase (middle), or C3 transferase washout (right) over the course of two hours. Increasing contractility with a C3 transferase washout produces significant cell shape changes. Scale bar is 10um.

Video 11: Light-mediated recruitment of prGEF causes irreversible junction shortening with HECD1 treatment (from Figure 4B). Zoomed in image of two Caco-2 cells during a 10 minute RhoA activation upon treatment with HECD1 and fluorescent secondary antibody. Junctions undergo similar junction length changes as WT cells. Scale bar is 10um.

Video 12: Light-mediated recruitment of prGEF causes internalization of HECD1-labeled E-cadherin puncta (From Figure 4B). Zoomed in image of a cell-cell junction undergoing a 10minute RhoA activation with HECD1 and fluorescent secondary antibody labelling. Labelled E-cadherin puncta compress, coalesce, and then are internalized within the slow phase of activation. Scale bar is 10um.

Video 13: Light-mediated recruitment of prGEF causes reversible junction shortening with SMIFH2 treatment (From Figure 4C, D). Zoomed in image of two Caco-2 cells during a 20 minute RhoA activation upon treatment with SMIFH2. Junctions contract with similar fast phases as WT, but the slow phase is prevented such that the junction returns to its original length upon light inactivation. Scale bar is 10um.

Video 14: Dynamin-mediated endocytosis regulates junction length changes (from Figure 5G). Zoomed in image of two Caco-2 cells, treated with Dynasore, during a 10 minute RhoA activation. The junction contracts but does not undergo junctional remodeling so it is not stabilized at a shorter length. The junction then returns to its original length upon RhoA removal. Scale bar is 10um.

Video 15: Periods of variable tension are necessary for increased junction length changes (from Figure 6E). Zoomed in image of Caco-2 cells undergoing two 20minute RhoA activations with a 20minute relaxation period in between. This type of cyclical activation, or ratcheting, produces marked junction length changes compared to one 40minute activation. Scale bar is 10um.

