## Supplementary figures and images for "RhoA mediates epithelial cell shape changes via mechanosensitive endocytosis"

### Supplemental Figures

Supplementary Figure 1

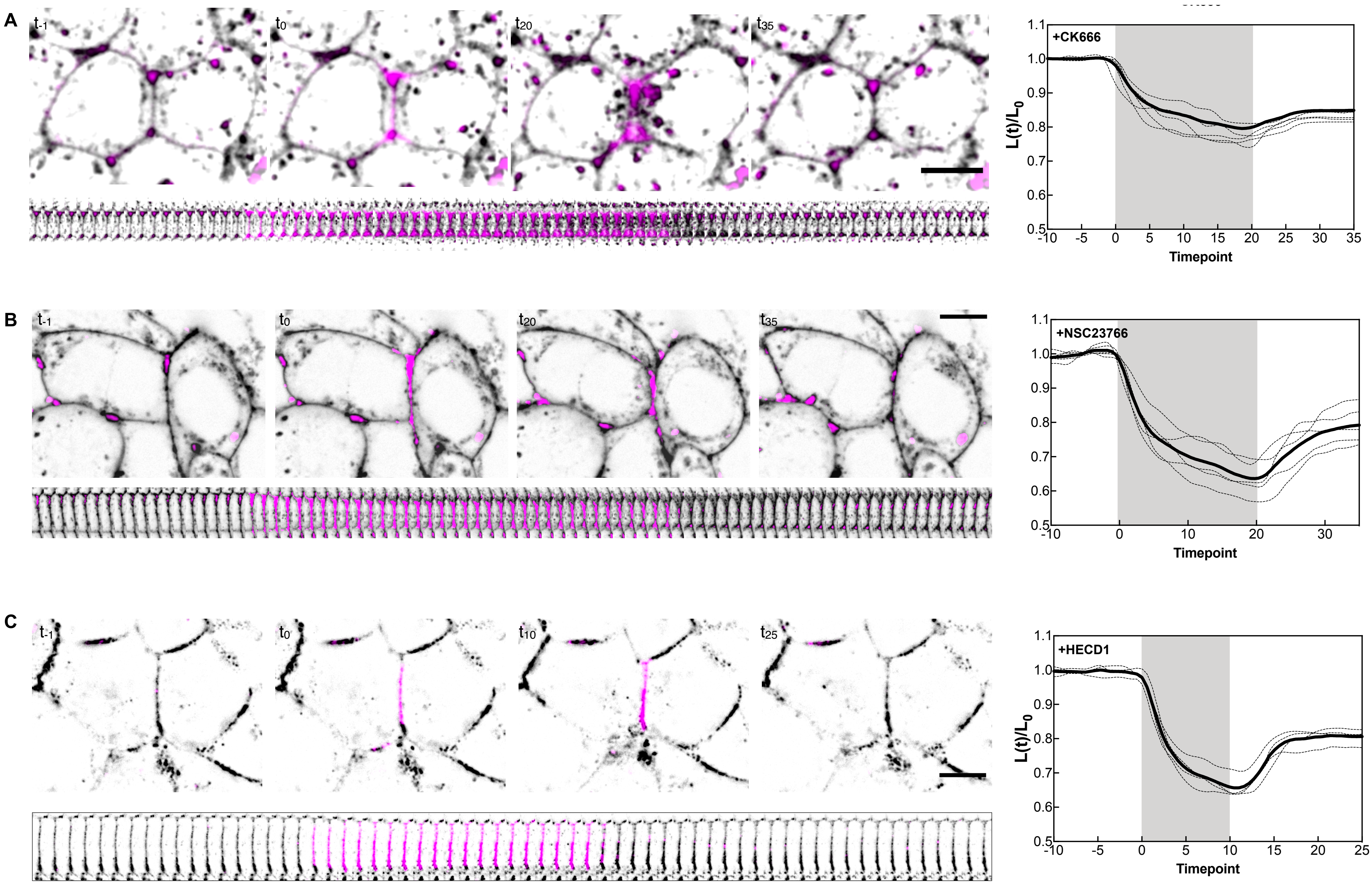

Supplementary Figure 2

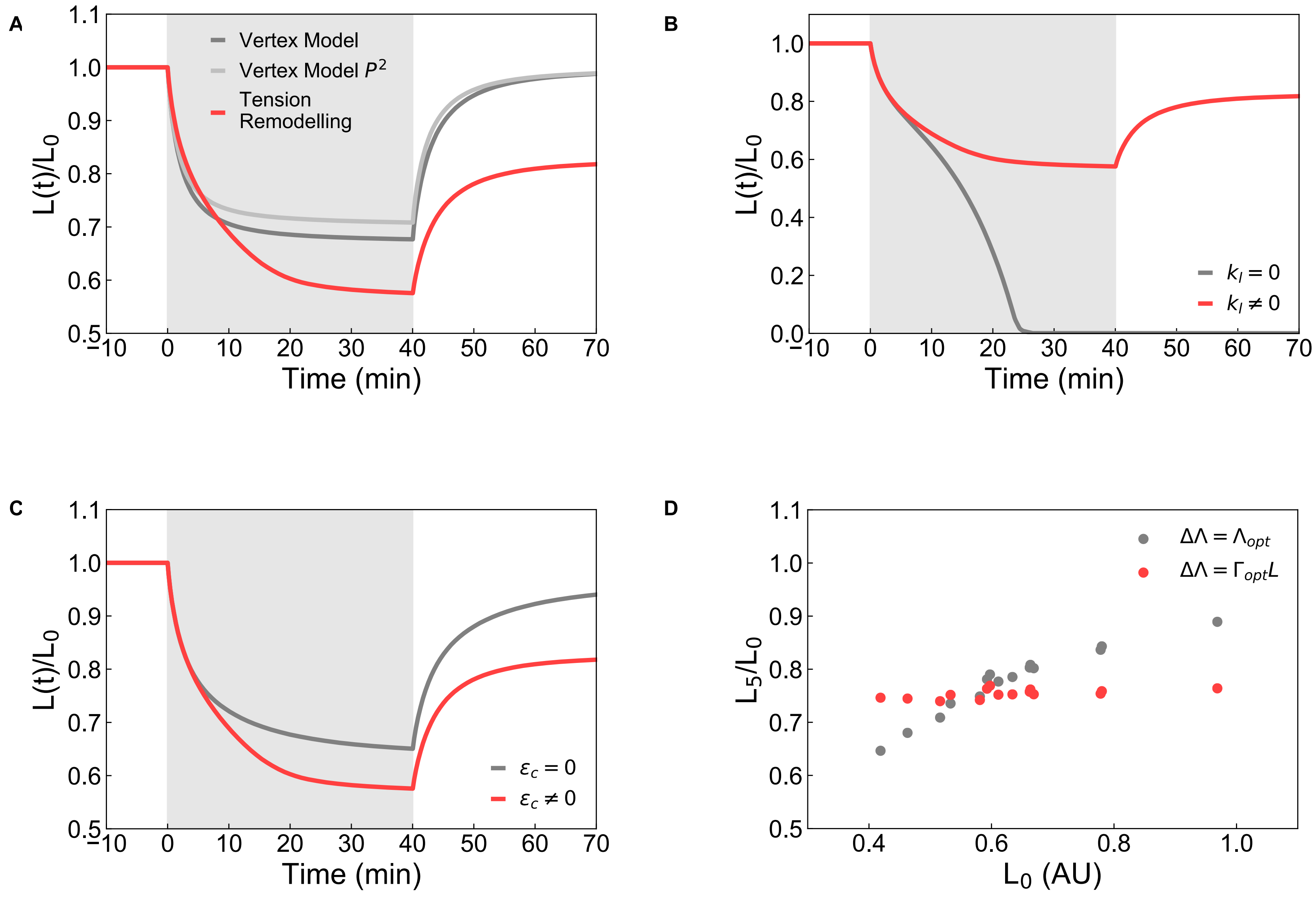

Supplementary Figure 4

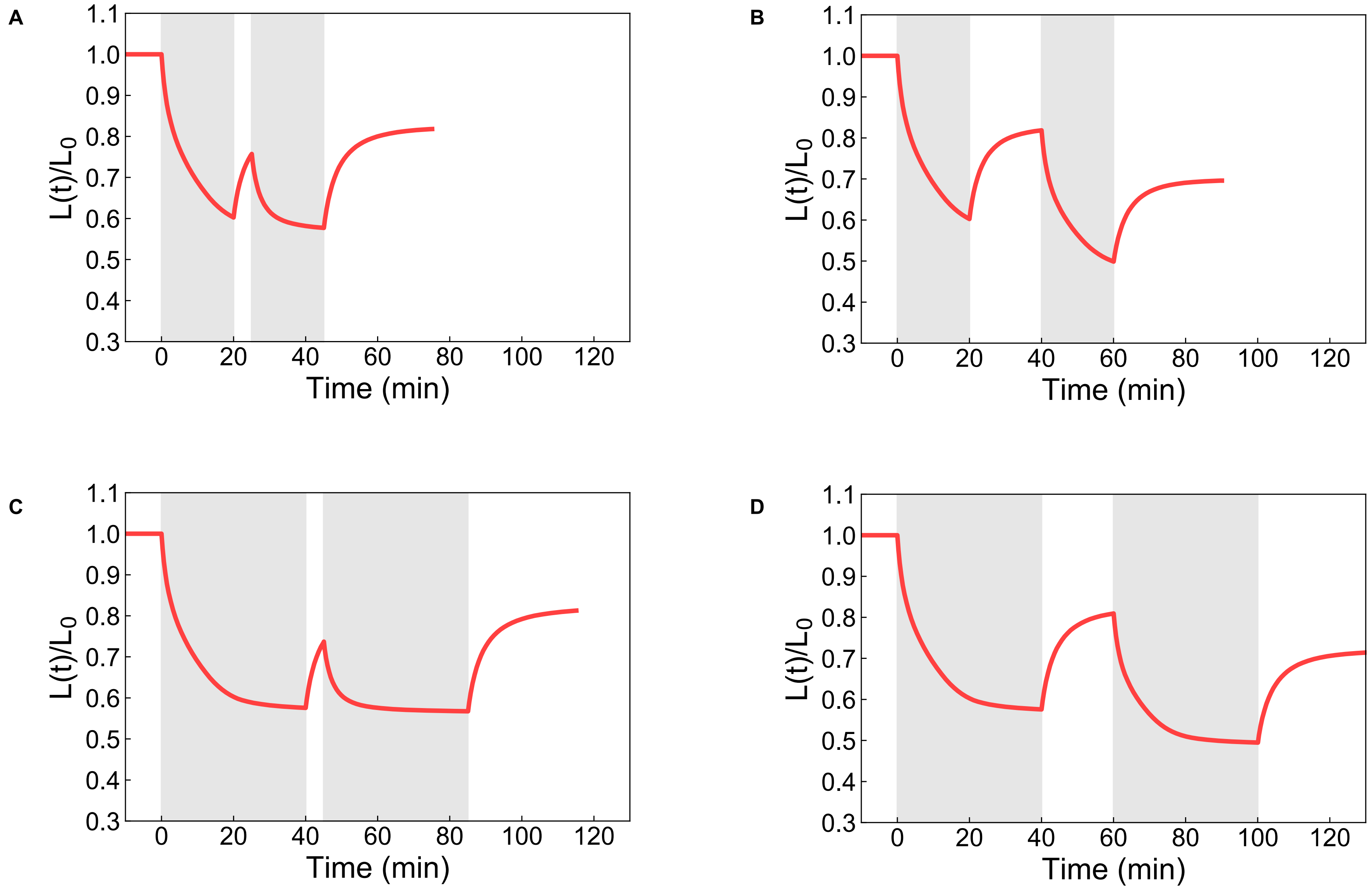
